# UTILIZATION OF A PUBLICLY AVAILABLE DIVERSITY PANEL IN GENOMIC PREDICTION OF *FUSARIUM* HEAD BLIGHT RESISTANCE TRAITS IN WHEAT

**DOI:** 10.1101/2023.02.15.528729

**Authors:** Z. J. Winn, J. H. Lyerly, G. Brown-Guedira, J. P. Murphy, R. E. Mason

**Affiliations:** North Carolina State University, Crop and Soil Sciences, Raleigh, North Carolina, United States of America; United States Department of Agriculture, Agricultural Research Service, Raleigh, North Carolina, United States of America; Colorado State University, Soil and Crop Sciences, Fort Collins, Colorado, United States of America

## Abstract

*Fusarium* head blight (FHB) is an economically and environmentally concerning disease of wheat (*Triticum aestivum* L). A two-pronged approach of marker assisted selection (MAS) coupled with genomic selection (GS) has been suggested when breeding for FHB resistance. An historical dataset comprised of entries in the Southern Uniform Winter Wheat Scab Nursery (SUWWSN) from 2011-2021 was partitioned and used in genomic prediction. Two traits were curated from 2011-2021 in the SUWWSN: percent *Fusarium* damaged kernels (FDK) and Deoxynivalenol (DON) content. Heritability was estimated for each trait-by-environment combination. A consistent set of check lines was drawn from each year in the SUWWSN, and K-means clustering was performed across environments to assign environments into clusters. Two clusters were identified for FDK and three for DON. Cross-validation on SUWWSN data from 2011-2019 indicated no outperforming training population in comparison to the combined dataset. Forward validation for FDK on the SUWWSN 2020 and 2021 data indicated a predictive accuracy *r* ≈ 0.58 and *r* ≈ 0.53, respectively. Forward validation for DON indicated a predictive accuracy of *r* ≈ 0.57 and *r* ≈ 0.45, respectively. Forward validation using environments in cluster one for FDK indicated a predictive accuracy of *r* ≈ 0.65 and *r* ≈ 0.60, respectively. Forward validation using environments in cluster one for DON indicated a predictive accuracy of *r* ≈ 0.67 and *r* ≈ 0.60, respectively. These results indicated that selecting environments based on check performance may produce higher forward prediction accuracies. This work may be used as a model to create a public resource for genomic prediction of FHB resistance traits across public wheat breeding programs.

**CORE IDEAS:**

1. The data from the Southern Uniform Winter Wheat Nursery may be used for genomic prediction.
2. Creating training populations based on like-check performance improves forward genomic predictive accuracies.
3. Filtering out locations with low genomic, per-plot, narrow-sense heritability may improve predictive accuracies.

## INTRODUCTION

*Fusarium* head blight (FHB), or head scab, is a fungal disease of wheat (*Triticum aestivum* L) and other small grains. In the United States, head scab is predominantly caused by the pathogen *Fusarium graminearum* (Ward et al., 2008). Disease development is favored by warm and wet conditions during flowering (Xu et al., 2007; Cowger et al., 2009). Infection can result in significant reduction in grain yield and quality, and subsequent mycotoxin accumulation creates additional food safety concerns (Buerstmayr et al., 2020; Ghimire et al., 2020; Ma et al., 2020)

Controlling FHB is challenging due to the complexity of the host-pathogen interaction. Chemical control has a limited window of application and resistance is quantitatively inherited (Buerstmayr et al., 2020). Breeding resistant cultivars is part of an integrated approach to mitigate losses from FHB, along with chemical control and cultural practices to reduce inoculum (Dill-Macky and Jones, 2000; Dweba et al., 2017; Shah et al., 2018). *Fusarium* species also produce the mycotoxin deoxynivalenol (DON), which is a vomitoxin that is harmful to human and animal health (Sobrova et al., 2010). For this reason, the United States Food and Drug Administration has set recommendations for the amount of DON allowable in wheat grain, and exceeding the threshold results in dockages when selling grain (McMullen et al., 1997).

Marker-assisted selection (MAS) has been used extensively to improve resistance to FHB and over 500 QTL with varying effect size have been identified (Dweba et al., 2017; Buerstmayr et al., 2020; Ghimire et al., 2020). While MAS is well suited for QTL with large effects, it is more challenging to implement for multiple smaller effect QTL (Heffner et al., 2009). Additionally, mapping populations frequently utilized in MAS studies to identify and validate QTL may not always represent the total germplasm in a breeding program or reflect performance in target environments. QTL stability in different genetic backgrounds and QTL by environment interactions are factors influencing both the choice of QTL to target and the potential for success in using MAS to incorporate those QTL in breeding programs (Xu and Crouch, 2008; Hospital, 2009). Utilizing genomic selection (GS) in conjunction with MAS may be beneficial for capturing these effects and furthering breeding for FHB resistance.

Genomic selection is the process of utilizing genome-wide markers and phenotypic data to predict phenotypic values for new breeding material (Meuwissen et al., 2001). In recent years, GS has been widely used for improvement of *Fusarium* head blight resistance and other agronomic traits in wheat (Rutkoski et al., 2011, 2014; Arruda et al., 2015; Lozada et al., 2019; Sandhu et al., 2021; Tomar et al., 2021; Gaire et al., 2022; Juliana et al., 2022). Achieving high prediction accuracies depends on various parameters relating to the quality of input data.

The genetic relationships between the individuals in the training set and the individuals in the prediction set, the quality of genotype data, and the quality of phenotype data all contribute to success of the training population. Prior studies have assessed strategies for identifying the most effective training populations, examining factors such as population size and structure, marker density, and composition based on genetic relationships or phenotypic distribution (Dawson et al., 2013; Arruda et al., 2015; Norman et al., 2018; Lozada et al., 2019; Verges et al., 2020).

The Southern Uniform Winter Wheat Scab Nursery (SUWWSN) is an annual collaborative nursery to evaluate resistance to FHB in elite germplasm under high disease pressure conditions. Entries from both public and private breeding programs are evaluated in mist-irrigated, inoculated nurseries and data is collected on multiple traits, including percent *Fusarium* damaged kernels (FDK) and DON content (Murphy et al., 2019, 2020). Annual reports are available at (https://scabusa.org/research-reportspublications, Accessed 2022-Aug-29). Collections of historical data, such as the SUWWSN, are valuable assets for genomic-assisted breeding.

Historical data sets from collaborative nurseries are a rich resource for constructing training populations for GS (Dawson et al., 2013; Sarinelli et al., 2019; Verges et al., 2020). Working with these curated data sets can present several challenges. In historical data sets, meta-information about individual trials or factors affecting trial environments may be limited due to changes in personnel, variation in data collection methods, or adoption of new technology over time. Historical data are often statistically unbalanced, with different breeding material being evaluated in each year. Successful implementation of GS relies on selecting informative data. For example, excluding low performing or poorly predictive environments from training populations can lead to increases in prediction accuracy (Heslot et al., 2013).

Clustering methods, such as principal components, k-means, or hierarchical clustering have been used in GS studies to account for population structure and assign genotypes to training populations based on genetic information (Norman et al., 2018; Andrade et al., 2019; Berro et al., 2019). Clustering can also be used to define mega-environments (Crespo-Herrera et al., 2021; Krause et al., 2022), which can inform environment selection for training populations and potentially improve predictions in target environments. Improvement in prediction accuracy depends on whether the mega-environments are distinct enough to justify using a clustering method versus using all available data (Dawson et al., 2013). Cluster analysis requires a complete data set where genotypes used to calculate a distance matrix are evaluated in each environment. In historical data sets, including the SUWWSN, the same check varieties are included in the nursery across years and locations. Analysis of check varieties can provide a basis for selecting year-location combinations that are informative for a training data set.

Heritability of a trait in an environment may be important when improving predictive accuracy. Training population subgroups partitioned based on heritability generated different prediction accuracies in maize populations, where the subgroups with higher heritability showed an increase in accuracy for grain yield, anthesis date, and plant height (Zhang et al., 2022). Prediction accuracy has generally been shown to increase with higher heritability (Jannink et al., 2010; Guo et al., 2014; Lozada et al., 2019), however this is dependent on the trait and the number of markers (Combs and Bernardo, 2013; Kaler et al., 2022).

The objectives of the presented work were to (i) investigate the use of the multi-environment SUWWSN historical data for prediction of percent FDK and DON accumulation, (ii) examine the impact of partitioning environments for training populations based on check performance on prediction accuracy for the FHB resistance traits FDK and DON in the SUWWSN, and (iii) investigate the use of partitioning environments for training populations based on heritability and the impact on prediction accuracy.

## MATERIALS AND METHODS

### Historical Data Curation

The phenotypic data used in this study is similar to the data set used by Winn et al (2022). Adjusted means of FDK and DON content in parts per million were compiled from the Southern Uniform Winter Wheat Scab Nursery (SUWWSN) reports from 2011-2021 (Murphy et al., 2015; Murphy & Navarro, 2010, 2011, 2012, 2013, 2014; Murphy et al., 2016, 2017, 2018, 2019, 2020, 2021). For the purposes of this study, only environments which had adjusted means reported for both FDK and DON were used for cross and forward validation. An environment, for this study, will refer to a specific location in a year.

Sampling the environments which reported adjusted means for both FDK and DON resulted in 45 environments from the years 2011-2019 and 15 environments from the years 2020-2021 for a total of 60 environments. Environments sampled include Arkansas, Louisiana, Virginia, Kentucky, North Carolina, Illinois, Missouri, South Carolina, and Texas (Figure 1). Across years 2011-2021, there were a total of 395 estimable genotypic values for both FDK and DON. Data was partitioned to separate years 2011-2019 from 2020-2021. All environments in the years 2011-2019 served as members of potential training populations and environments in the years 2020-2021 were used for forward validation.

**Figure 1.**
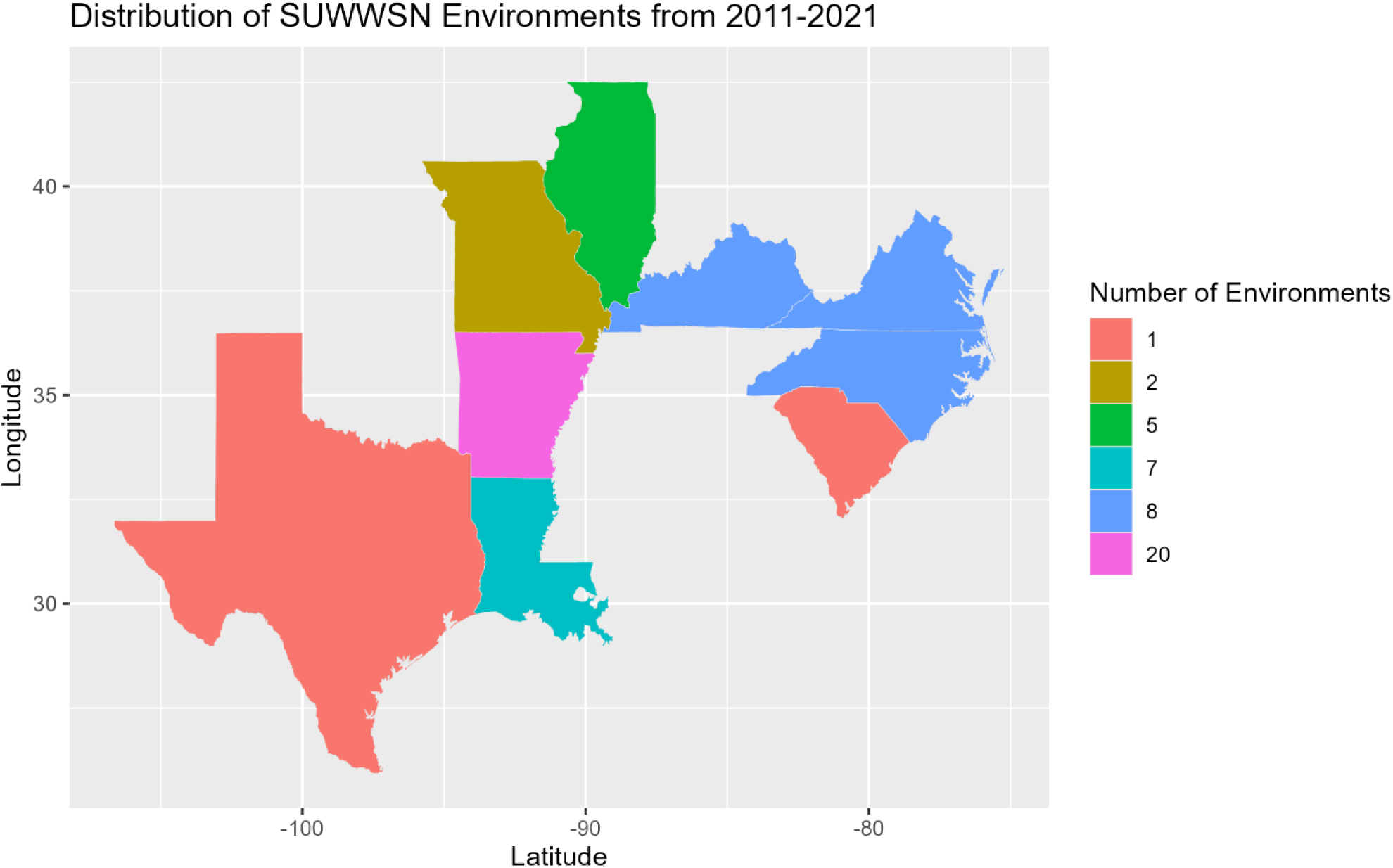
A visualization of total number of environments in the 2011-2021 Southern Uniform Winter Wheat Scab Nursery. Displayed is a graph with latitude on the x axis and longitude on the y axis. Each state shown above is filled according to the number of environments contributed by that state to the dataset, and the number is indicated by the legend to the right of the graph.

### Nursery Organization and Phenotyping

Each environment reported in the SUWWSN was grown in some incarnation of a corn-spawn inoculated and mist irrigated nursery specific to each program. For examples of a corn-spawn inoculated and misted FHB nursery conducted by North Carolina State University, refer to methods in Peterson et al (2016, 2017). Four checks were replicated across all years of the SUWWSN: three considered moderately resistant and one highly susceptible.

Of the four repeated checks in the SUWWSN, ‘Jamestown’ (PI 653731), is a moderately FHB resistant line developed by Virginia Polytechnic Institute and State University (Griffey et al., 2010). Jamestown has had multiple FHB resistance QTL identified in its haplotype (Carpenter et al., 2020; Wright, 2014). ‘Bess’ (PI 642794) is an FHB moderately-resistant line developed by the Missouri Agricultural Experiment Station (McKendry et al., 2007). Bess has also had several FHB resistance QTL identified in its haplotype (Petersen et al., 2016, 2017). ‘Ernie’ is a moderately FHB resistant line developed by the Missouri Agricultural Experiment Station, which has had several FHB resistance QTL identified in its haplotype (Brown-Guedira et al., 2008; McKendry et al., 1995). ‘Coker 9835’ is an FHB susceptible line developed by Northup, King, and Company (AR, United States) (Subramanian et al., 2016).

Percent *Fusarium* damaged kernels was determined by referencing known standards of scabby seed in set increments and providing a visual estimate of the percentage of seeds in a sub sample which appeared affected. Seeds were considered affected if they had a chalky, white-to-pink appearance. Deoxynivalenol content was measured by gas chromatography via collaborating laboratories in the United States Wheat and Barley Scab Initiative network. Protocols for sampling, submission, and estimation of DON in seed samples can be found at the United States Wheat and Barley Scab Initiative web domain [https://www.scabusa.org/don_labs]. All other phenotyping protocols can be found at the same web domain [https://www.scabusa.org]. Adjusted means of FDK and DON were calculated on a collaborator-by-collaborator basis and reported via the SUWWSN reports.

### Genotyping

Genotyping protocols in this study were as written in Winn et al (2022). Leaf tissue at the four-leaf stage was taken for each genotype in the SUWWSN panel and DNA was extracted using sbeadex plant maxi kits (LGC Genomics, Middlesex, UK) as directed by the manufacturer’s protocol. Genotyping-by-sequencing was performed according to Poland et al (2012). Libraries were sequenced at 192 plex densities and processed on an Illumina HiSeq 2500. SNP discovery using raw data was done via the Tassel-5GBSv2 pipeline version 5.2.35 (Glaubitz et al., 2014).

Reads were aligned to the RefSeq 1.0 wheat genome assembly (Appels et al., 2018) using the Burrows-Wheeler aligner version 0.7.12 (Li & Durbin, 2009). Data was filtered by removing genotypes with 85% or more missing data, SNPs with a minor allele frequency of 5% or lower, SNPs which had a heterozygous call frequency of 10% or higher, SNPs missing 20% or more data, SNPs with average read-depth of less than 1 or more than 100 and reads that aligned to unknown chromosomes. Imputation via Beagle 5.2 was conducted post filtering (B. L. Browning et al., 2018; S. R. Browning & Browning, 2007).

### Data Analysis

All data analysis was performed using R statistical software version 4.2.1 (R Core Team, 2013). Adjusted means of FDK and DON were checked for violation of linearity and normality by visual assessment of their respective distributions. All mixed linear models were conducted using the “asreml()” function from the ASREML-R package in R (Butler et al., 2009). Best linear unbiased estimates (BLUEs) were calculated for genotypes using adjusted means by environment via the following mixed linear model:

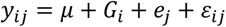

Where *y* is the response, *μ* is the mean, *G* is the fixed genotype effect, *e* is the random environment effect 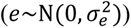, and *ε* is the residual error 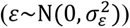. Genotypic variance was estimated in each trait-by-environment combination by the following genomic best linear unbiased prediction (gBLUP) model:

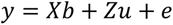

Where y is the response, X is the design matrix for fixed effects, b is the vector of fixed effects, Z is the design matrix for random genetic effects, and u is the vector of additive genetic effects defined by the genomic relationship matrix (VanRaden, 2008).

Per-plot, narrow-sense, genomic heritability for each trait-by-environment combination of the SUWWSN was calculated using variances derived from gBLUP models:

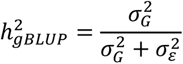

Where 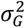 is the genotypic variance and 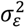 is the error variance. The estimated genomic, narrow-sense, per-plot heritability was used to select the environment-by-trait combinations that would be evaluated in potential training populations.

Adjusted means for the nursery wide checks (Jamestown, Bess, Ernie, and Coker 9835) that span across the years 2011-2019 were compiled into a matrix of n genotypes by m environments, where each cell represented the adjusted mean of a check for either FDK or DON in an environment. These matrices were passed to the function “NbClust()” from the package “NbClust”(Charrad et al., 2014). The “NbClust()” function calculates distance matrices from data sets and provides 30 different clustering indexes to indicate the optimal number of clusters in a dataset. For this study, the “index” argument “all” was indicated, and a subset of 26 different clustering algorithms were considered. These indexes included: “KL”, “CH”, “Hartigan”, “CCC”, “Scott”, “Marriot”, “TrCovW”, “TraceW”, “Friedman”, “Rubin”, “Cindex”, “DB”, “Silhouette”, “Duda”, “Pseudot2”, “Beale”, “Ratkowsky”, “Ball”, “Ptbiserial”, “Frey”, “McClain”, “Dunn”, “Hubert”, “Sdindex”, “Dindex”, and “SDbw” (Charrad et al., 2014). Clusters were used to indicate possible training populations. From the possible 26 indexes calculated for the dataset, a majority rule vote is taken, and the most frequent number of clusters indicated as optimal is considered the optimal clustering overall.

For both FDK and DON check values across 2011-2019, the number of possible clusters entertained ranged from 2 to 15 and the optimal clustering of the data was decided by majority rule across all available indexes. A principal component analysis (PCA) was conducted on the matrices to visualize the optimal clustering indicated by the “NbClust()” function on the first two principal components (PCs).

For both cross and forward validation, genomic estimated breeding values (GEBVs) were generated by calculating best linear unbiased predictions (BLUPs) for individuals in the test population using the gBLUP model:

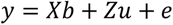

Where y is a vector of responses (where missing values are maintained for individuals in the test population), X is the design matrix for fixed effects, b is the vector of fixed effects, Z is the design matrix for random genetic effects, and u is the vector of additive genetic effects defined by the genomic relationship matrix (VanRaden, 2008). Cross validated accuracy distributions were obtained from 100 permutations of a cross-validation scheme where 80% of the data within a training population was randomly kept in as training and the remaining 20% was left out as a test population.

Significant differences of training populations under study vs the combined training population utilizing all the environments were calculated as a function of Z-scores. The Z-score used to estimate significant differences from the combined population was calculated as follows:

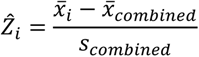

Where 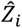 is the calculated Z-score of training population 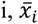 is the estimated mean of the distribution for training population 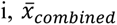 is the estimated mean of the combined training population, and *s_i_* is the estimated standard deviation of the combined training population. Values where 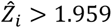 or 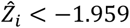 were considered significantly different.

### Prediction Procedure – Training Population Nomenclature and Data Organization

The SUWWSN dataset was partitioned into a training set including all the environments from 2011-2019 and a validation set including environments from 2020 (SUWWSN20) and 2021 (SUWWSN21). Several methods of filtering were assessed for the training data. Heritability 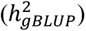 was used to filter environments in the training set for FDK prediction and for DON prediction. Heritability thresholds entertained for filtering out environments were as follows:

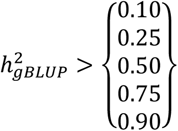

Environments were retained in the respective training population if their 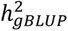 was greater than the values on the right-hand side of the above equation. For training populations created through filtering the data via heritability, the population name will be denoted as “*h2 > x trait*,” where x denotes the value of 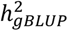 used to filter data and trait represents either FDK or DON.

Clustering of environments based on check performance was conducted and clusters were used to denote different training populations. Clusters were visualized on a PCA graph of the first PC versus the second PC. Training populations which were made through clustering will be denoted as “*cluster x trait*,” where x denotes the cluster and trait represents either FDK or DON. Predictions made with the total available dataset are denoted as “*combined trait”*, where trait represents either FDK or DON.

## RESULTS AND DISCUSSION

### Data Assessment – Clustering of Environments and Estimation of Heritabilities

Of the 26 methods of clustering tested for FDK, 24 of the indexes produced definitive results. Clustering among the indexes indicated the following number of clusters: nine indexes proposed two clusters, three proposed three clusters, five proposed four clusters, three proposed six clusters, one proposed eleven clusters, one proposed 14 clusters, and two proposed 15 clusters. By majority vote, the number of clusters identified as optimal for FDK was two.

Of the 26 methods of clustering tested for DON, 23 of the indexes produced definitive results. Clustering among the indexes indicated the following number of clusters: five indexes proposed two clusters, six proposed three, two proposed four, one proposed six, one proposed nine, one proposed 12, one proposed 14, and four proposed 15. By majority vote, the number of clusters identified for DON was three.

A principal component analysis was performed on the matrices for FDK and DON check values and the indicated clusters by the “NbClust()” algorithm were visualized (Figure 2, 3). For FDK, the first two PCs captured approximately 32% and 17% of the total variation, respectively. For DON, the first two PCs captured approximately 12% and 7% of the total variation, respectively.

**Figure 2.**
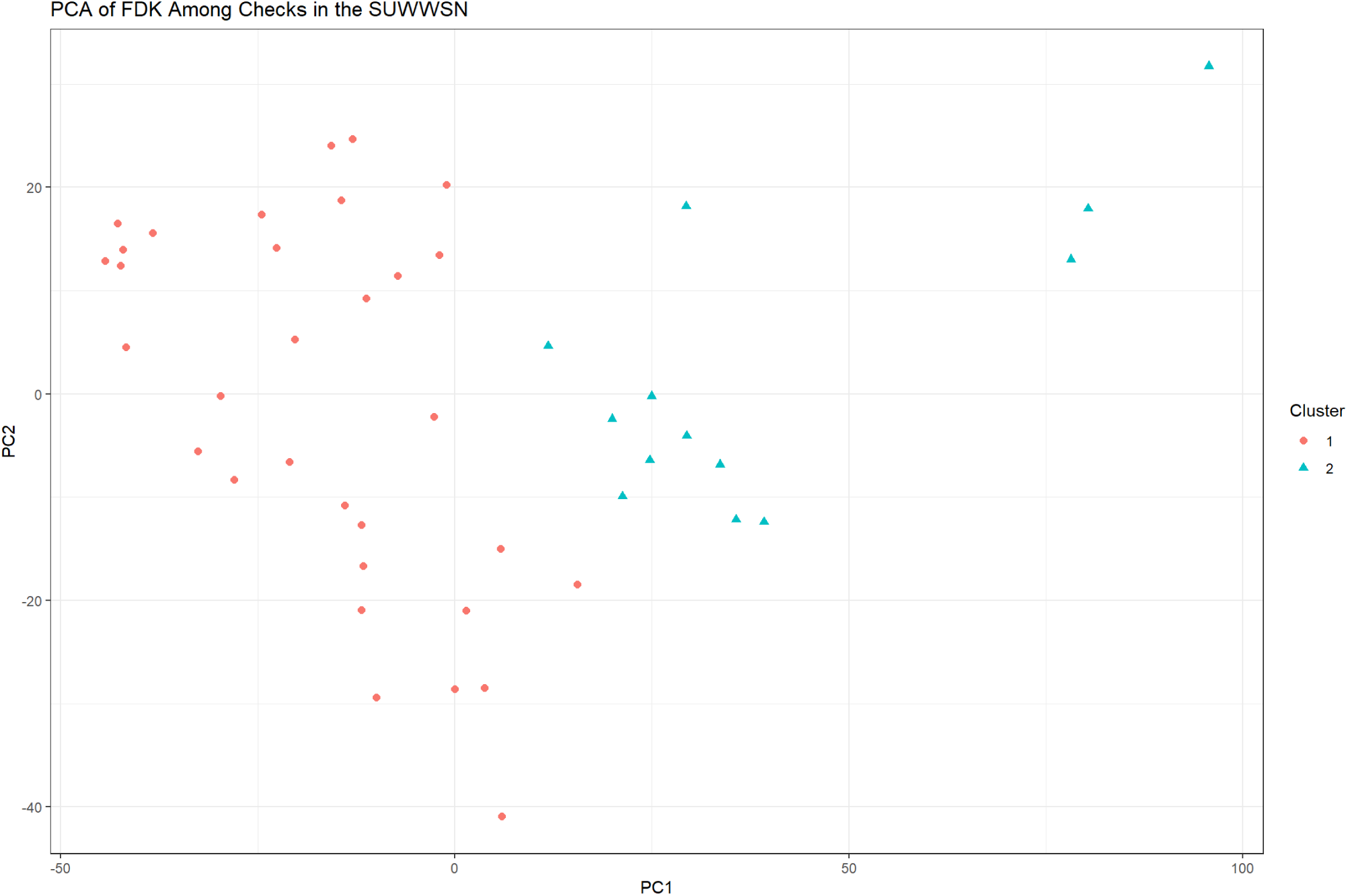
A comparison of principal component one (PC1) vs. principal component two (PC2) derived from a principal component analysis (PCA) of percent *Fusarium* damaged kernels (FDK) in the 2011-2019 southern uniform winter wheat scab nursery (SUWWSN). Clusters indicated by majority rule vote from the “NbClust()” function are color and shape coded and defined by the legend on the righthand side of the graph.

**Figure 3.**
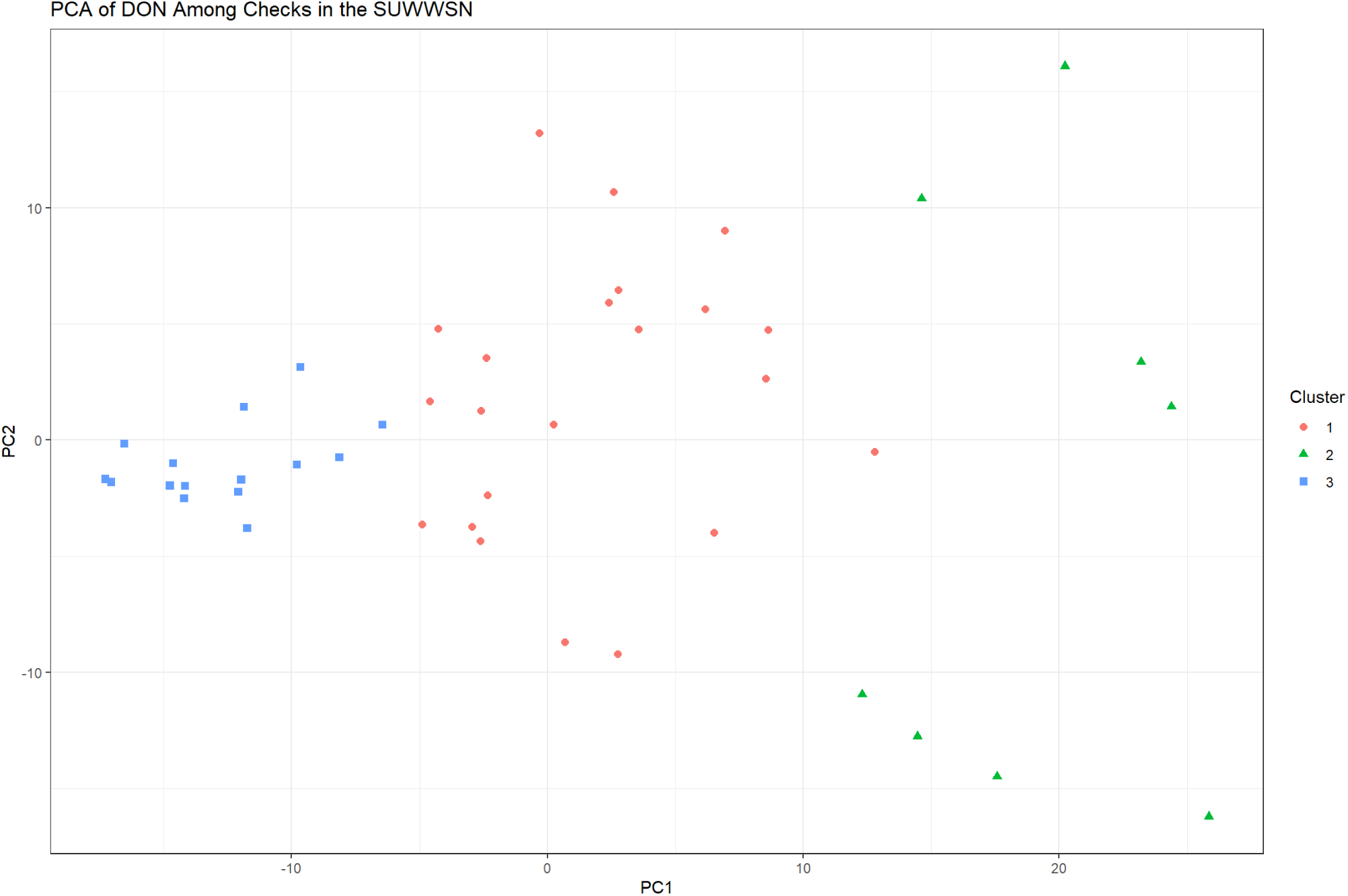
A comparison of principal component one (PC1) vs. principal component two (PC2) derived from a principal component analysis (PCA) of deoxynivalenol content (DON) in the 2011-2019 southern uniform winter wheat scab nursery (SUWWSN). Clusters indicated by majority rule vote from the “NbClust()” function are color and shape coded and defined by the legend on the righthand side of the graph.

Estimated heritability for each environment-by-trait combination, calculated via gBLUP, and the cluster to which each environment-by-trait combination in belongs is provided (Table 1). Training population sizes were partitioned based on cluster and heritability (Table 2). To understand the difference in check averages between training populations, the BLUEs of check values from each training population were visually compared (Figure 3, 4).

**Table 1.**
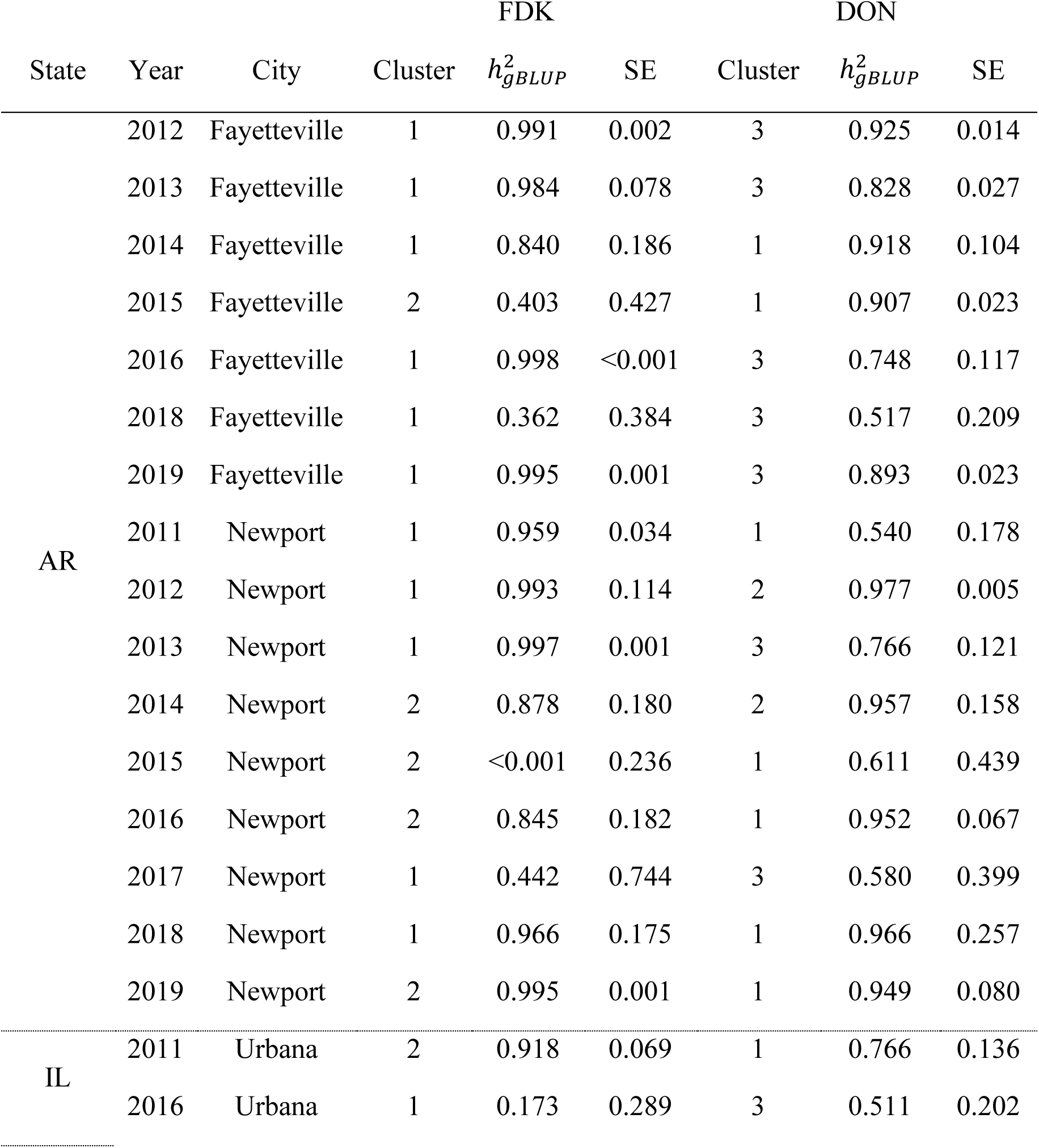
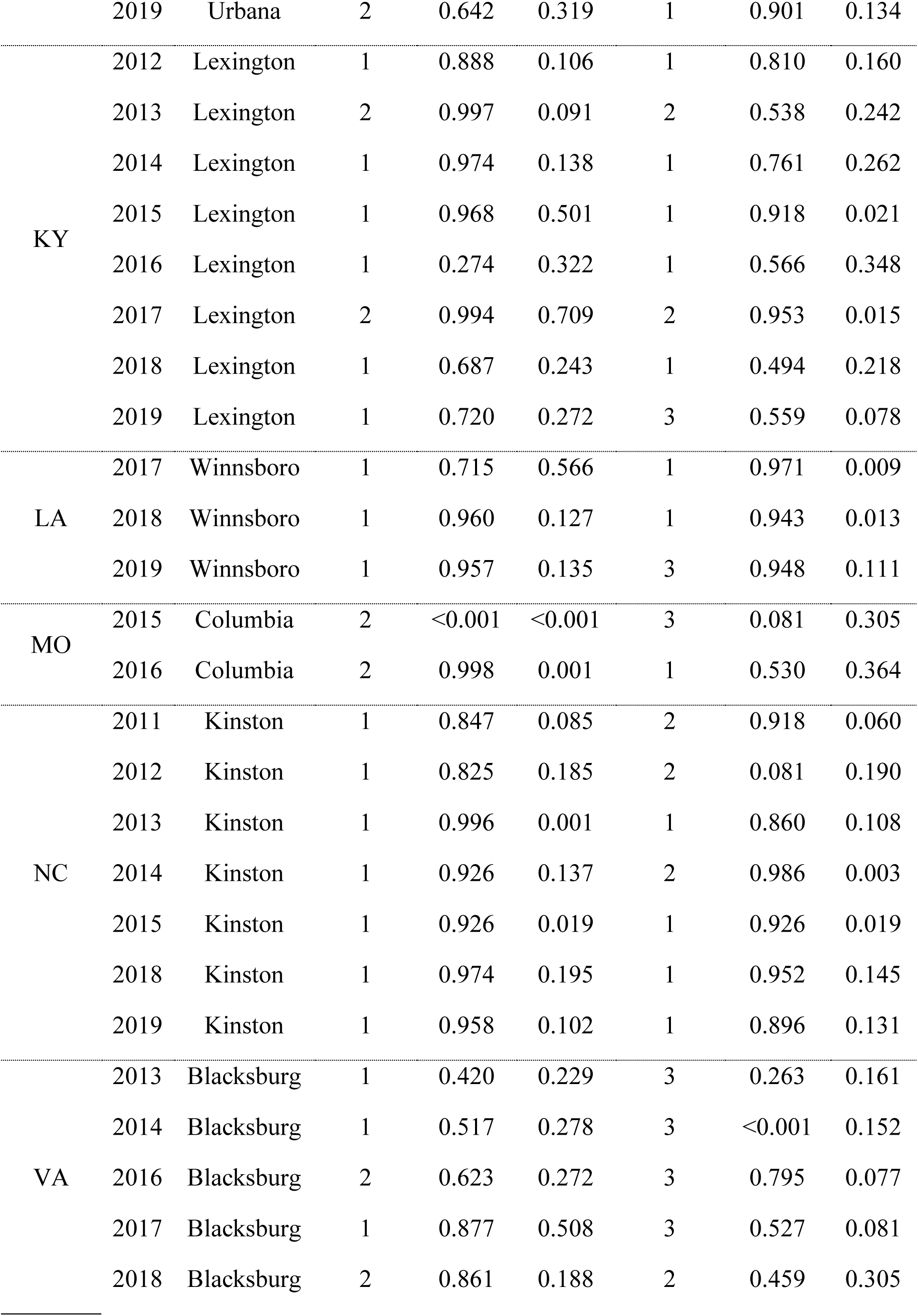
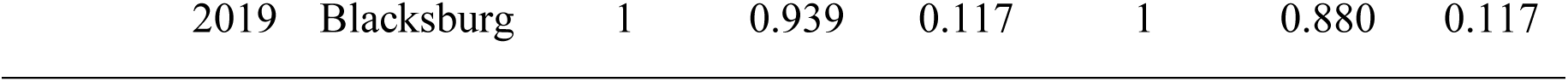
List of each location by state, year, and city from the 2011-2019 Southern Uniform Winter Wheat Scab Nursery. Columns under Fusarium damaged kernels (FDK) or Deoxynivalenol (DON) indicate what cluster each environment groups with based on check performance, what the heritability of that trait in that environment is, and the standard error of the heritability associated with that trait in that environment. Values less than 0.001 are denoted as “<0.001”.

**Table 2.** A list of each training population tested and the total number of observations, genotypes, and environments in each training population.

| Training Population | Observations | Genotypes | Environments |
| --- | --- | --- | --- |
| *Combined FDK* | 1847 | 331 | 45 |
| *Cluster 1 FDK* | 1354 | 331 | 32 |
| *Cluster 2 FDK* | 493 | 297 | 13 |
| *h2>0.10 FDK* | 1791 | 331 | 43 |
| *h2>0.25 FDK* | 1756 | 331 | 42 |
| *h2>0.50 FDK* | 1574 | 331 | 37 |
| *h2>0.75 FDK* | 1357 | 331 | 31 |
| *h2>0.90 FDK* | 1001 | 328 | 23 |
| *Combined DON* | 1847 | 331 | 45 |
| *Cluster 1 DON* | 871 | 331 | 22 |
| *Cluster 2 DON* | 374 | 244 | 8 |
| *Cluster 3 DON* | 602 | 277 | 15 |
| *h2>0.10 DON* | 1721 | 331 | 42 |
| *h2>0.25 DON* | 1721 | 331 | 42 |
| *h2>0.50 DON* | 1586 | 331 | 39 |
| *h2>0.75 DON* | 1180 | 331 | 28 |
| *h2>0.90 DON* | 694 | 283 | 18 |

**Figure 4.**
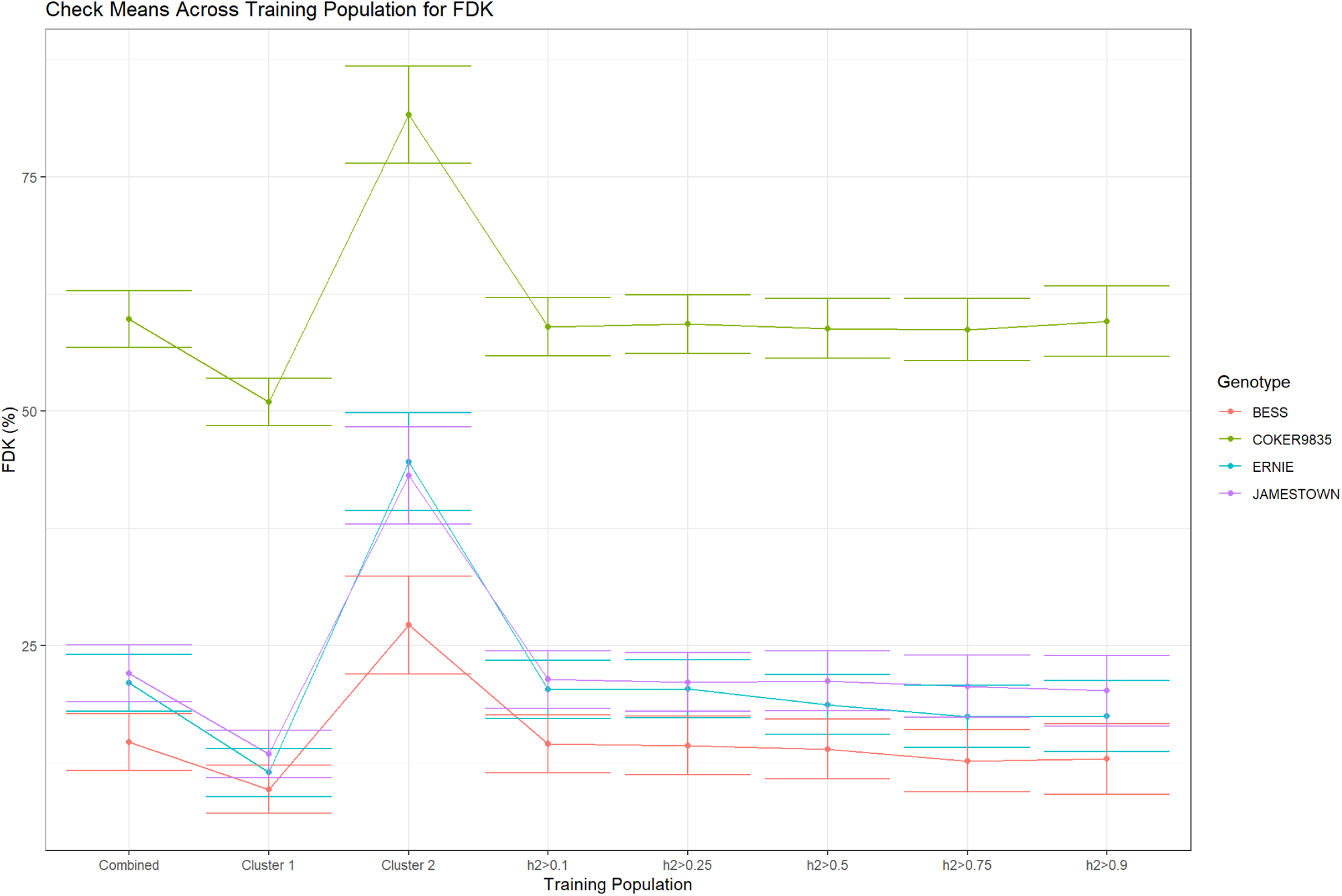
Best linear unbiased estimates (BLUEs) for check varieties from each training population for percent *Fusarium* damaged kernels (FDK). Each color represents a different genotype shown in the figure legend on the righthand side of the graph. Value of FDK in percent visual scabby seeds is labeled on the y axis, and the x axis shows what training population the BLUEs belong to. Bars surrounding point estimates of BLUEs represent one standard error about the estimate.

**Figure 5.**
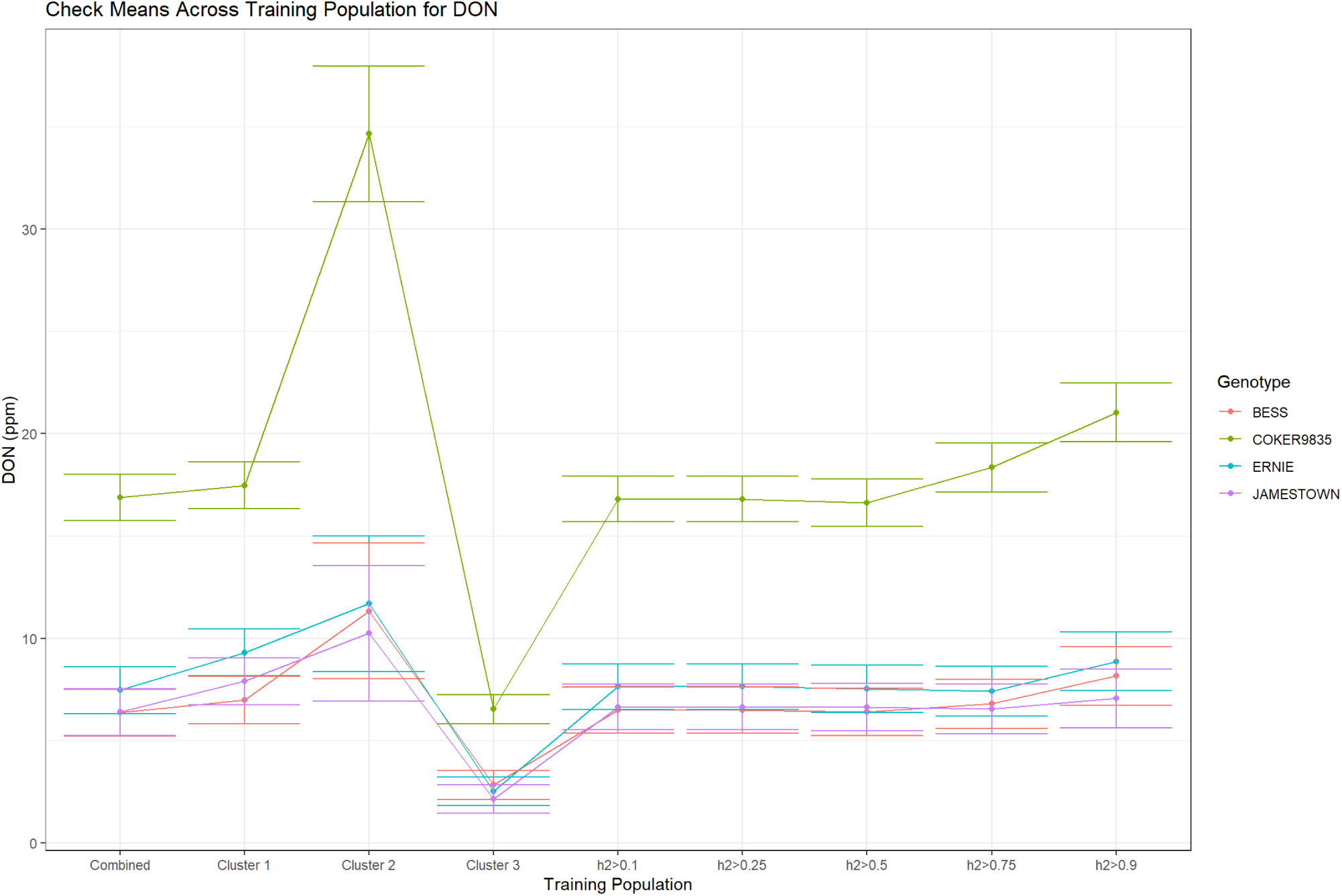
Best linear unbiased estimates (BLUEs) for check varieties from each training population for deoxynivalenol (DON) content in parts per million (ppm). Each color represents a different genotype shown in the figure legend on the righthand side of the graph. Value of DON in ppm is labeled on the y axis, and the x axis shows what training population the BLUEs belong to. Bars surrounding point estimates of BLUEs represent one standard error about the estimate.

Check BLUEs tended to vary more dramatically between clusters than between different subsets of environments based on heritability. This is expected, because clusters were made based on similarity between check performance, and heritability is not directly related to performance of checks within an environment.

Furthermore, no discernable pattern between clustering based on check performance and subsetting based on heritability was evident in the dataset. Partitioning based on like-check performance often resulted in differing number of estimable genotypic values, while partitioning based on heritability tended to make no major change in the number of estimable genotypic values. A reduction in the number of estimable genotypes was observed for training populations partitioned via high heritability 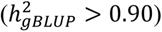.

*Cluster 1 FDK* had check BLUEs which did not vary one standard error outside the *Combined FDK* BLUEs; however, *Cluster 2 FDK* had check BLUEs which obviously deviated from the *Combined FDK* BLUEs. In *Cluster 2 FDK*, checks tended to have higher estimates of FDK than either the *Combined FDK* BLUEs or the *Cluster 1 FDK* BLUEs.

*Cluster 1 DON* appeared to have check BLUEs which were within one standard error of the *Combined DON* BLUEs. *Cluster 2 DON* appeared to have BLUEs which were substantially higher than either *Combined DON* or *Cluster 1 DON. Cluster 3 DON* appeared to have the lowest BLUEs for all checks among the training populations for DON.

These trends of differing check mean values in clusters for FDK and DON may explain the distance between environment PCs observed in the PCAs, and thus, average check mean values within environment is the main contributing factor to explain PC1 and PC2 for both FDK and DON. This may be further extrapolated as the “disease pressure” within environments, where high disease pressure environments result in higher FDK and DON check values, while lower disease pressure environments result in lower FDK and DON check values. Therefore, the clustering of environments based on check values may be grouping environments with like disease pressure.

### Cross and Forward Validation Accuracies

Cross validation yielded no meaningful differences in predictive accuracies for any of the different training populations for either FDK or DON (Figure 6, 7). Prediction accuracies among *Combined FDK, Cluster 1 FDK, Cluster 2 FDK, h2 > 0.10 FDK, h2 > 0.25 FDK, h2 > 0.50 FDK, h2 > 0.75 FDK,* and *h2 > 0.90 FDK* appeared similar with overlapping distributions, except in the case of *Cluster 2 FDK* which appeared to have a visibly lower (yet not statistically lower) distribution mean (Table 3). Prediction accuracies among *Combined DON, Cluster 1 DON, Cluster 2 DON, Cluster 3 DON, h2 > 0.10 DON, h2 > 0.25 DON, h2 > 0.50 DON, h2 > 0.75 DON,* and *h2 > 0.90 DON* also appeared relatively similar with overlapping distributions. Like the case of *Cluster 2 FDK*, *Cluster 2 DON* appeared to have a visibly lower (yet insignificantly lower) distribution mean than the rest of the training populations.

**Figure 6.**
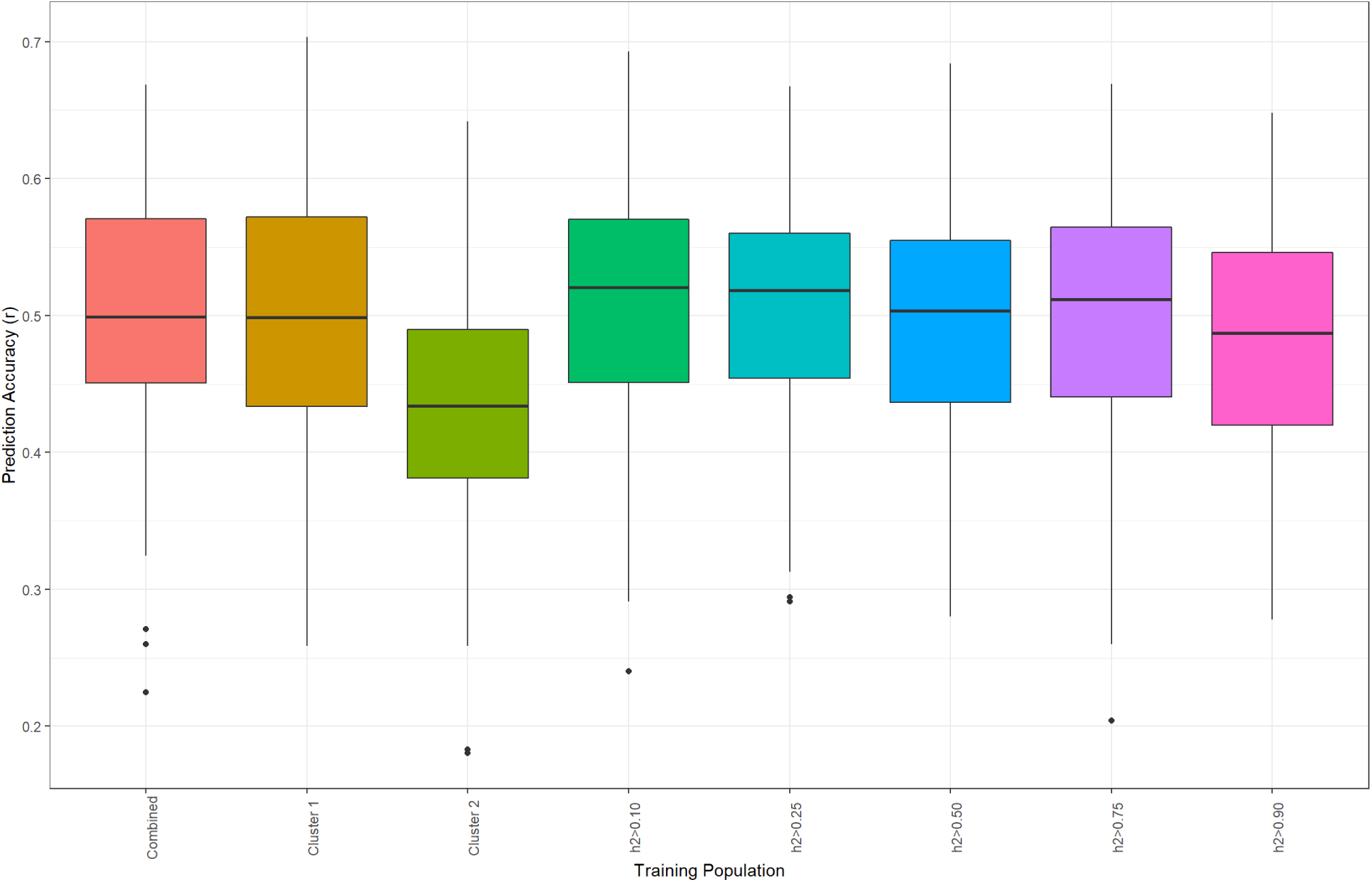
Graphical display of 100 permutations of cross-validation accuracies for all different training populations for *Fusarium* damaged kernels (FDK). The y axis displays the Pearson’s correlation coefficient (or prediction accuracy) between the genomic best linear unbiased prediction of the testing population and the best linear unbiased estimate of the testing population. The x axis displays the name of each training population tested.

**Figure 7.**
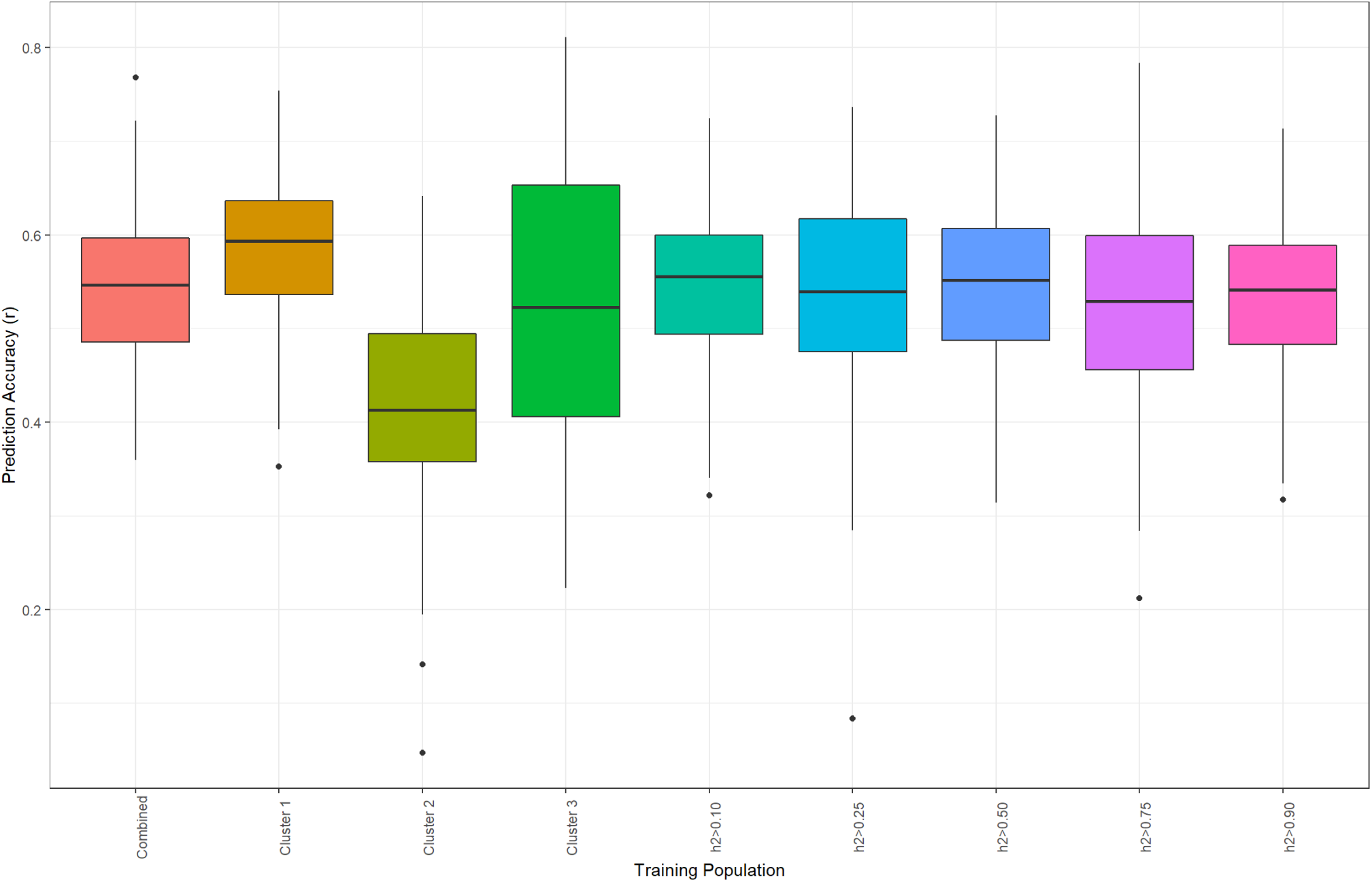
Graphical display of 100 permutations of cross-validation accuracies for all different training populations for deoxynivalenol content (DON). The y axis displays the Pearson’s correlation coefficient (or prediction accuracy) between the genomic best linear unbiased prediction of the testing population and the best linear unbiased estimate of the testing population. The x axis displays the name of each training population tested.

**Table 3.** Summarization of cross-validation results. The resulting minimum, maximum, mean, standard deviation (SD), and Z-score calculated in reference to the combined training population’s distribution is listed for each training population of *Fusarium* damaged kernels (FDK) and deoxynivalenol content (DON).

| Trait | Training Population | Min | Max | Mean | SD | Z-score |
| --- | --- | --- | --- | --- | --- | --- |
| FDK | *Combined* | 0.22 | 0.67 | 0.50 | 0.09 | - |
|  | *Cluster 1* | 0.26 | 0.70 | 0.50 | 0.09 | -0.03 |
|  | *Cluster 2* | 0.18 | 0.64 | 0.43 | 0.08 | -0.81 |
|  | *h2>0.10* | 0.24 | 0.69 | 0.51 | 0.09 | 0.15 |
|  | *h2>0.25* | 0.29 | 0.67 | 0.51 | 0.09 | 0.08 |
|  | *h2>0.50* | 0.28 | 0.68 | 0.50 | 0.09 | -0.02 |
|  | *h2>0.75* | 0.20 | 0.67 | 0.49 | 0.09 | -0.06 |
|  | *h2>0.90* | 0.28 | 0.65 | 0.48 | 0.09 | -0.18 |
| DON | *Combined* | 0.36 | 0.77 | 0.54 | 0.08 | - |
|  | *Cluster 1* | 0.35 | 0.75 | 0.58 | 0.08 | 0.52 |
|  | *Cluster 2* | 0.05 | 0.64 | 0.42 | 0.11 | -1.06 |
|  | *Cluster 3* | 0.22 | 0.81 | 0.52 | 0.15 | -0.10 |
|  | *h2>0.10* | 0.32 | 0.72 | 0.55 | 0.08 | 0.11 |
|  | *h2>0.25* | 0.08 | 0.74 | 0.53 | 0.11 | -0.05 |
|  | *h2>0.50* | 0.31 | 0.73 | 0.55 | 0.09 | 0.10 |
|  | *h2>0.75* | 0.21 | 0.78 | 0.52 | 0.10 | -0.15 |
|  | *h2>0.90* | 0.32 | 0.71 | 0.53 | 0.08 | -0.09 |

For forward validation in the SUWWSN20 and SUWWSN21 for FDK, higher predictive accuracies than the *Combined FDK* training population were observed in the *Cluster 1 FDK* training population. In the SUWWSN20 and SUWWSN21 forward validation schemes, prediction accuracies for *Cluster 1 FDK* were 8% and 5% higher (respectively) than the *Combined FDK* training population’s prediction accuracies. Likewise for DON, prediction accuracies for *Cluster 1 DON* were consistently higher for SUWWSN20 and SUWWSN21 in comparison to the *Combined DON* training population. In the SUWWSN20 and SUWWSN21 forward validation schemes, prediction accuracies for *Cluster 1 DON* were 10% and 9% higher (respectively) than the *Combined DON* training population’s prediction accuracies (Table 4).

**Table 4.**
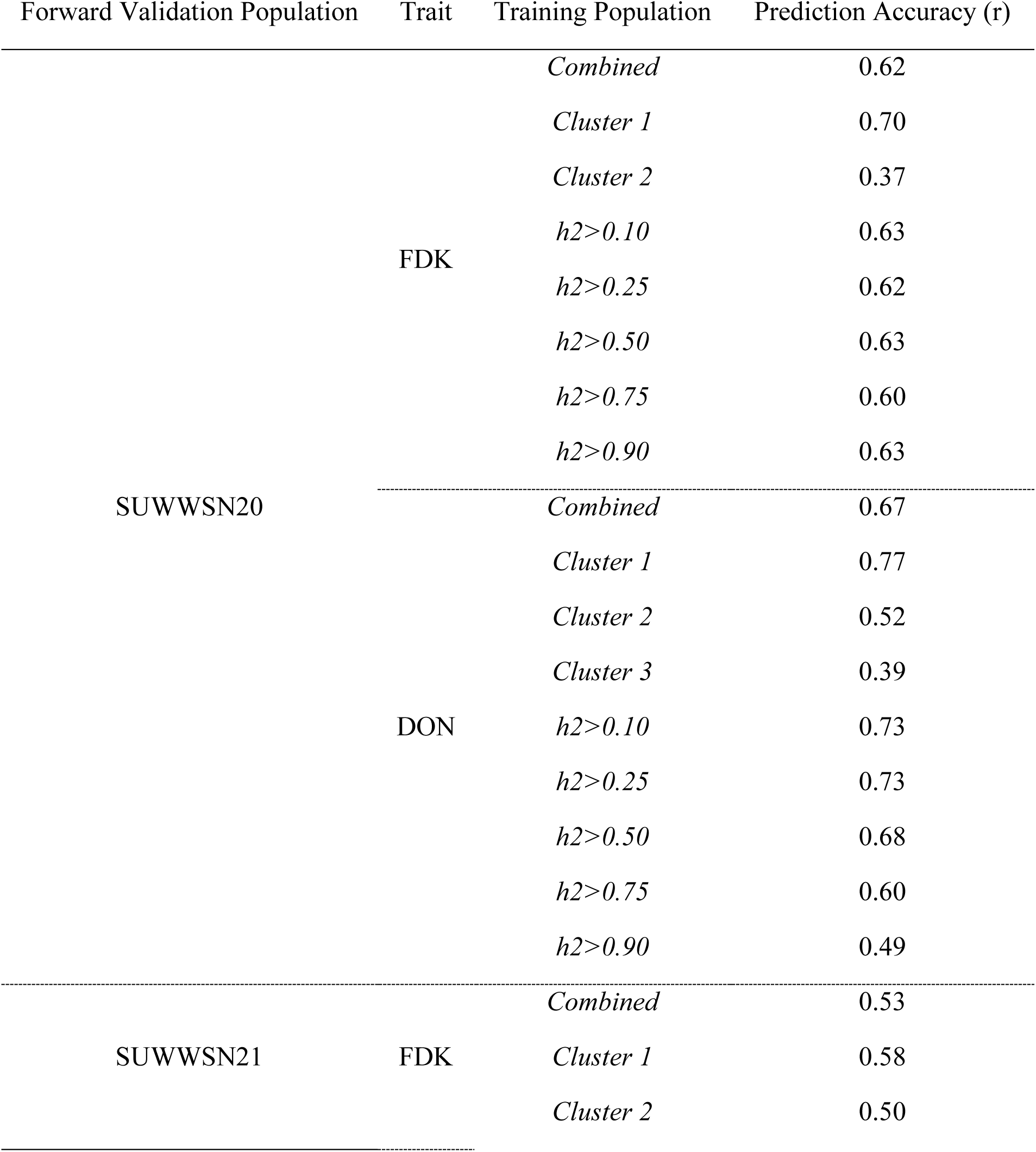
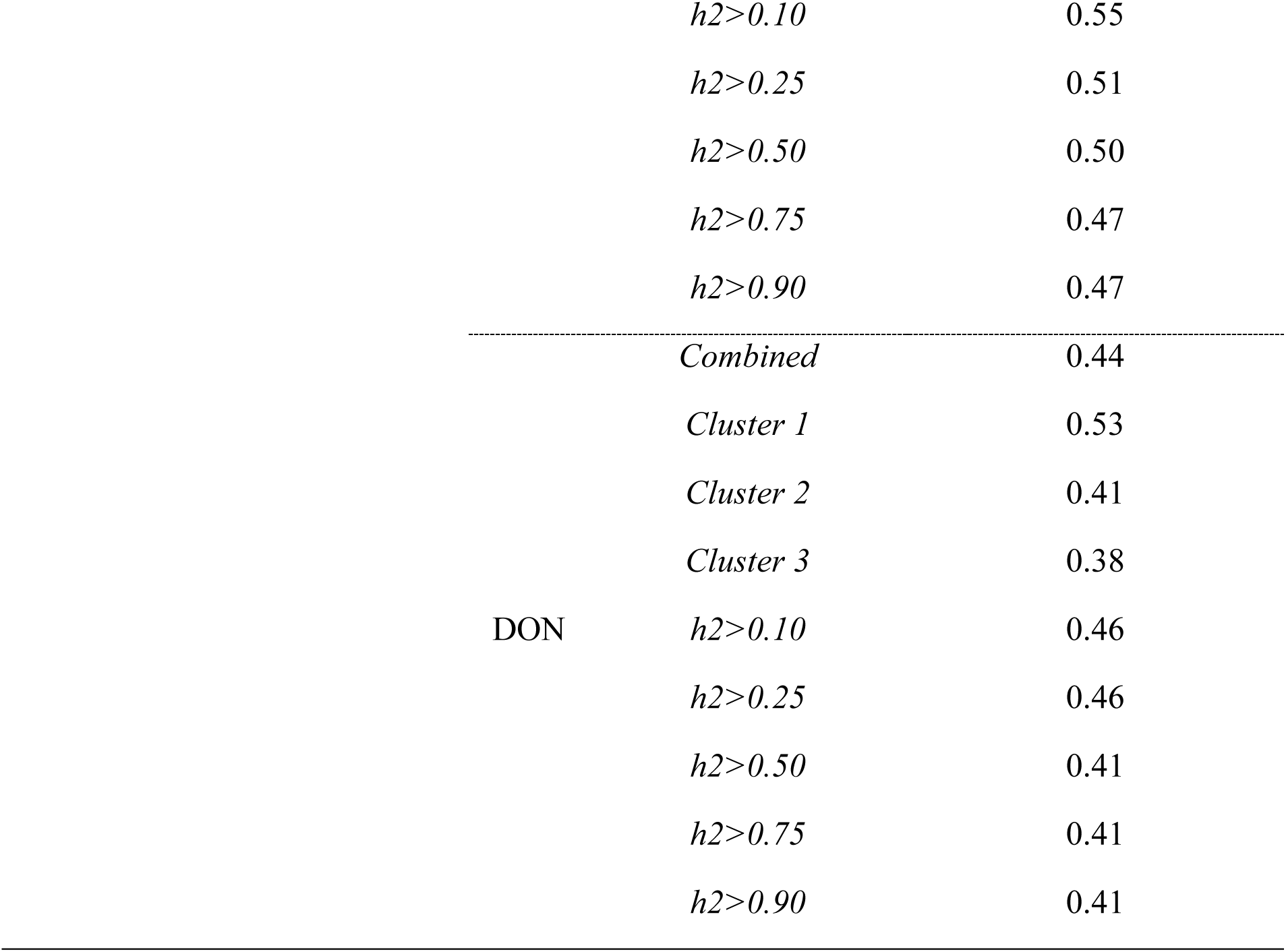
Summary table of forward validation prediction accuracies in the Southern Uniform Winter Wheat Scab Nursery (SUWWSN) for the 2020 (SUWWSN20) and 2021 (SUWWSN21) years for multi-environment BLUEs of percent Fusarium damaged kernels (FDK) and deoxynivalenol content (DON).

Clusters beyond 1 for either FDK or DON either performed equivalently to the combined dataset or substantially worse than the combined dataset for either trait. In the SUWWSN20 and SUWWSN21 forward validation scheme for FDK, *Cluster 2 FDK* performed 25% and 3% worse in terms of prediction accuracy than the *Combined FDK* training set. In the SUWWSN20 and SUWWSN21 forward validation scheme for DON, *Cluster 2 DON* performed 15% and 3% worse than *Combined DON*, and *Cluster 3 DON* performed 28% and 6% worse than *Combined DON*. *Cluster 1 FDK* and *Cluster 1 DON* were superior to *Combined FDK* and *Combined DON* in prediction accuracy, meaning that there is potential benefit in partitioning environments based on check values in prediction of FHB reaction traits.

In both the SUWWSN20 and SUWWSN21 forward validation schemes, partitioning environments based on heritability for either FDK or DON resulted in prediction accuracies that were comparable to or moderately worse (1-5% worse) than the combined training sets for either trait. One exception to this pattern is forward validated prediction accuracy for DON in SUWWSN20 in respect to using the *h2>0.10 DON* and *h2>0.25 DON* training populations. In SUWWSN20, *h2>0.10 DON* and *h2>0.25 DON* produced prediction accuracies 5% higher than the *Combined DON* training population. However, this trend was not consistent among years. Therefore, partitioning environments by heritability in this dataset may not be of as high utility as previously reported for other traits in wheat.

## DISCUSSION

Selecting representative environments with high quality data to include in training populations is a key component in achieving high prediction accuracy in GS. Historical data from collaborative nurseries can serve as potential resources for breeding programs and prior studies in wheat have demonstrated the utility of historical data sets as a basis for training populations (Sarinelli et al., 2019; Verges et al., 2020). In the work presented, historical data from the SUWWSN was used to generate training populations to predict FHB disease resistance traits. We investigated whether partitioning based on trait heritability or clusters made by like-check performance resulted in improved prediction accuracy.

Multiple studies have demonstrated that accounting for the relationships among genotypes and increasing the size of the training population results in higher prediction accuracy (Berro et al., 2019; Norman et al., 2018). However, fewer studies have examined the impact of selecting environments based on quality parameters. In this study, we have demonstrated that partitioning environments by like-check performance may substantially increase forward validated prediction accuracies for both FDK and DON content.

Prior studies in maize (*Zea maize* L.) have shown that prediction accuracy may increase when using high heritability environments to predict traits of interest (Zhang 2022). In this study, the distributions of training populations created by partitioning via heritability were not significantly different from the combined training population in cross validation.

In forward validation, prediction accuracy decreased for DON as partitioning environments by heritability became very restrictive. In forward validation of the SUWWSN20, prediction accuracy for DON decreased from 0.67 using the *Combined DON* training population to 0.49 when using the *h2>0.90 DON* training population. SUWWSN21 accuracy for DON decreased from 0.44 with the *Combined DON* training population to 0.41 when using the *h2>0.90 DON* training population. For FDK there was no noted difference in prediction accuracy with training populations partitioned based on heritability in forward validation in SUWWSN20 (*Combined FDK*=0.62 and *h2>0.90 FDK*= 0.63). In forward validation of FDK in the SUWWSN21, there was a reduction in accuracy when stringently partitioning environments via heritability (*Combined FDK*=0.53 and *h2>0.90 FDK*= 0.47).

For DON, the *h2>0.90 DON* training population contained 18 environments, compared to 45 environments using the *Combined DON* training population. Studies have shown that including more environments, and more diversity, can improve prediction accuracy (Norman et al., 2018). This reduction in the number of environments may not capture the range of disease severity or genetic diversity, since not all lines are tested in each environment.

These results indicated that severely limiting the number of environments by using only the high heritability environments may reduce predictive ability in the data set. However, training populations partitioned based on removing low heritability environments 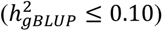 did produce a small increase in forward validation prediction accuracy for FDK and DON. By these results, it may be recommended to remove very low heritability environments 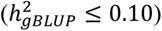 from training populations.

Cluster analysis of environmental check means generated two training populations for FDK, (*Cluster 1 FDK* and *Cluster 2 FDK*) and three training populations for DON (*Cluster 1 DON*, *Cluster 2 DON*, and *Cluster 3 DON*). Visual comparison of BLUEs across cluster illustrated that the mean values of check varieties were different between moderately resistant and susceptible varieties.

The *Cluster 3 DON* training population was the lowest performing training population in forward validation. The *Cluster 3 DON* training population contained 277 estimable genotypic values in 15 environments and had the smallest separation between the BLUEs among moderately resistant and susceptible checks. The *Cluster 3 DON* training population also had a wide distribution for cross validation accuracy, which indicated that considering low disease pressure environments as training data resulted in greater variation in accuracy for predicting DON, depending on the composition of the specific permutation.

In the case of the *Cluster 3 DON* training population, lower forward validation accuracies could be due to low disease pressure environments comprising much of this training population. It was shown that the BLUEs of check varieties in *Cluster 3 DON* were substantially lower than other training populations and that the separation between moderately resistant and susceptible varieties was narrower than other training populations. Multiple studies have demonstrated that the quality of input data is critical to the success of GS (Rutkoski et al., 2015; He et al., 2016; Hoffstetter et al., 2016; Belamkar et al., 2018) and using environments where check varieties are poorly distinguished is ineffective as a training population.

The *Cluster 2 DON* and *Cluster 2 FDK* training population did not outperform the *Combined* or *Cluster 1* training populations for predicting FDK or DON in forward validation. The *Cluster 2* training populations for FDK and DON had the lowest mean accuracy and distribution in cross validation for their respective traits. While the differences in accuracy with cross validation were not significant, a modest reduction in accuracy was observed. There was less overlap in the distribution of cross validation accuracy for DON between predictions made with the *Cluster 2 DON* training population and those of other training sets, which indicated that data partitioned into the *Cluster 2 DON* training population may not be as well suited for making predictions as other training populations.

The *Cluster 2* training populations contained 297 estimated genotypic values from 13 environments for FDK and 244 estimated genotypic values from eight environments for DON. For both traits, the *Cluster 2* training population was comprised of the smallest number of environments. Furthermore, the *Cluster 2* training population for either FDK or DON included high disease pressure environments, as illustrated in the BLUEs for the check varieties.

Previous research has indicated that removing outlier environments with atypical levels of biotic or abiotic stress can be beneficial for improving prediction accuracy (Michel et al., 2016). This must be balanced against reducing the number of genotypes represented in the training population, particularly when utilizing historical data sets where not all genotypes are represented in each year (Isidro et al., 2015). As with partitioning based on heritability, reducing the number of environments may also fail to account for the genetic diversity in germplasm or target environments, and therefore reduce predictive ability in the data set.

Results from this study show that *Cluster 1* training populations performed best in forward prediction for both FDK and DON. Forward prediction accuracy for FDK increased from 0.62 to 0.70 and from 0.53 to 0.58 compared to the *Combined FDK* training population for each year, respectively. For DON, accuracy increased from 0.67 to 0.77, and 0.44 to 0.53 compared to the *Combined DON* training population for each year, respectively.

The *Cluster 1* training populations were comprised of environments with a range of heritability for FDK and DON. The range of heritability was 0.49 – 0.97 for *Cluster 1 DON* and 0.17 – 0.99 for *Cluster 1 FDK*. The mean heritability for the *Cluster 1* training populations was 0.82 for DON and 0.81 for FDK. This was slightly higher than the mean for the *Combined* training population in each case, where the mean heritability was 0.73 in the *Combined DON* training population and 0.78 for *Combined FDK*.

The environments with low heritability 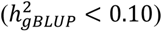 were not removed prior to clustering analysis. For FDK these low heritability environments were found in the *Combined FDK* and *Cluster 2 FDK* training populations. For DON, low heritability environments were partitioned into the *Combined DON*, *Cluster 2 DON*, and *Cluster 3 DON* training populations. While not directly selecting against low heritability environments, it appears that clustering based on check performance within environments did remove exceptionally low heritability environments from the *Cluster 1* training populations. This further reinforces the idea that removing very low heritability environments may be a best practice in training population design (Isidro et al., 2015; Michel et al., 2016).

For FDK and DON, the BLUEs for the check varieties for *Cluster 1* were most similar to those of their respective *Combined* training sets. For FDK, *Cluster 1* check variety BLUEs were slightly lower than the estimates from the *Combined* training population, however resistant and susceptible checks were still clearly distinguished. The number of estimated genotypic values for *Cluster 1 FDK* and *Cluster 1 DON* was 331, which is the same as that of *Combined FDK* and *Combined DON* training populations. Therefore, the number of estimated genotypic values did not change, despite the reduction in the number of observations in the training populations.

Historical data sets, including the SUWWSN, are valuable tools for genomic prediction. Here we have demonstrated that data from the SUWWSN can be used successfully to predict the performance of new breeding material. Best forward prediction accuracies for two years were 0.70 and 0.58 for FDK, and 0.77 and 0.53 for DON. From these results, we suggest that removing very low heritability environments (*h_gBLUP_* < 0.10), followed by clustering of like check performance, may be a best practice in partitioning data in historical datasets for use in training GS model.

## ACKNOWLEDGMENTS

Thank you to the United States Wheat and Barley Scab Initiative for providing the funding from the years 2010 to 2022 for the data used in this research. Thank you to all collaborators which participate in the annual screening of the Southern Uniform Winter Wheat Scab Nursery. Thank you to individuals which report and curate information for the Southern Uniform Winter Wheat Scab Nursery. This project was supported by funds derived from the North Carolina Small Grains Growers Association and the Competitive Grant 2022-68013-36439 (WheatCAP) from the USDA National Institute of Food and Agriculture

